# Behavioral assays to study neural development in Xenopus laevis tadpoles

**DOI:** 10.1101/2020.08.21.261669

**Authors:** Arseny S. Khakhalin, Virgilio Lopez, Carlos Aizenman

## Abstract

Escape responses, orienting reflexes, and social behaviors in Xenopus laevis tadpoles have been well documented in the literature (Lee et al. 2010; Roberts et al. 2000; Simmons et al. 2004; Katz et al. 1981; Villinger and Waldman 2012). In this article, we describe several behavioral protocols that together allow researchers efficiently (in terms of financial cost and time investment) and effectively assess developmental abnormalities in pre-metamorphic Xenopus tadpoles.

## 1. Schooling

Schooling behavior experiments allow for the observation of tadpole social interactions and in the past has been used as a method to characterize behavioral deficits in models of neurodevelopmental disorders (James et al., 2015). Unlike other species of frogs, Xenopus tadpoles show polarized schooling. Not only do tadpoles aggregate, they also swim in the same direction. Quantifying both aggregation and relative swim angle can give us an important measure of social behavior and sensory integration (Wassersug et al. 1981). Past iterations of these experiments have required the continued presence of an experimenter throughout the duration of each trial and relied upon expensive software for subsequent data analysis. The instrument configuration and analysis protocol outlined here provide an automated method to assess schooling by delivering a series of timed vibratory stimuli to a group of tadpoles to induce swimming behavior, and then controlling a camera (Iturbe 2020) to document their positions via still images. Both stimulus delivery and image acquisition are automated using the Python programming language. Analysis is done using ImageJ (Schindelin, et al. 2012) and custom Python scripts which are provided in this protocol. The specific equipment configuration, and scripts shown show here provide one solution, but other equipment and custom scripts can be substituted.

## Materials

### Reagents

- Xenopus rearing media (see **Appendix A** below for options and recipes)

### Equipment

- **LabJack U3-HV** – an inexpensive analog-digital data acquisition device for controlling vibration delivery by a computer
- **Jintai Dental Laboratory Vibrator** – to provide vibratory stimulus **GoPro Hero 7** (with any flexible mount of choice) – to acquire video and still images
- Any **LED light tracing tablet** (eg. Picture/Perfect light pad) – to provide ilumination
- **IOT Controllable Power Relay** – for powering vibrator, activated by the LabJack DAQ
- **17 cm diameter flat bottom glass bowl** – experimental arena

### Software

- Python 3.X: https://www.python.org/
- Jupyter: https://jupyter.org/install.html
- FIJI: https://imagej.net/Fiji/Downloads
- LabJack drivers and software: https://labjack.com/support/software/installers
- **LabJack Python library:** https://labjack.com/support/software/examples/ud/labjackpython
- Custom scripts for this project (from this repository, https://github.com/khakhalin/Xenopus-Behavior)

## Methods

### Software installation

1. **Install Python 3, and Jupyter notebooks. We recommend using Anaconda** distribution package, but installing Python and Jupyter separately would also work. Step-by-step instructions can be found here: https://jupyter.readthedocs.io/en/latest/install/notebook-classic.html
2. Download or clone the GitHub repository located at:https://github.com/khakhalin/Xenopus-Behavior For this protocol in particular, you will only need the files in the folder `01_Schooling`.
3. Follow the instructions in the instructions.md file to download and setup all necessary software and drivers to run the LabJack-U3 and GoPro.

### Equipment setup

1. Connect one wire from the FIO4 pin on the LabJack-U3, to the positive terminal on the IOT power relay’s removable screw terminal block and one wire from the adjacent GND pin on the LabJack-U3 to the negative terminal (**Fig. 1A**).
2. Plug the LabJack-U3 into your computer via USB. The green light on the device should illuminate.
3. Plug the dental vibrator into one of the “normally off” outlets on the Power Relay and ensure that the vibrator is turned on high.
4. Connect the power relay to an outlet and switch on, a red light should appear.
5. Open a Command prompt/Terminal window, ensure that you are in the directory with the supplied Python programs and run the following command: `$ Python u3vibrate.py` This should trigger a series of 4 vibrations, indicating that you have successfully installed the appropriate drivers to connect to the LabJack-U3.
6. Set up the camera above the arena using a flexible mount. Once the camera has been successfully connected to the computer via WIFI, open a second Command prompt/Terminal window and run the following command: `$ Python GoProStream.py` A live feed of your GoPro camera should open. You are now ready to run your experiment.
7. Secure the illumination pad to the dental vibrator platform with doube-sided tape or Velcro to provide illumination from the bottom.
8. Add 350 mL of 10% Steinberg’s Solution to a large round dish, 17cm in diameter and place the dish on the dental vibrator, directly beneath the GoPro (Note: It is helpful to include some barriers to hold the bowl in place, should the vibrations cause it to move throughout the duration of the experiment.) Use the screen on the GoPro to adjust the placement of the vibrator and dish to ensure it is in the camera’s frame. The distance between the GoPro and the arena will vary depending on the specifics of your setup, typically it is around 50 cm above the dish.
9. Add 15 to 20 stage 46-49 Xenopus laevis tadpoles to the dish and be sure to leave an object of known length somewhere in frame, a small ruler or some marked distance of known length.
10. Run the following command in the first Command Prompt window: `$ Python schooling_experiment.py`. The entire experiment is executed using a programmed loop. The loop consists of an image acquisition command, a 150 second wait time, a vibratory stimulus command, and another 150 second wait time. Therefore, each image and stimulus event is separated by a 300 second wait period, but the two events are offset from each other by 150 seconds. This loop is set to execute for 4000 seconds, which in the end will result in 12 acquired images for each experiment. Images acquired by the GoPro are saved as automatically named .JPG files, making the sequence clear. The timing parameters for the experiment can be easily altered by the experimenter by changing the values for the timer (line 29) and wait times (lines 46 & 50) in the script.

### Data analysis – FIJI

This section describes how to record position and orientation of tadpoles in each frame.

**Figure 1.**
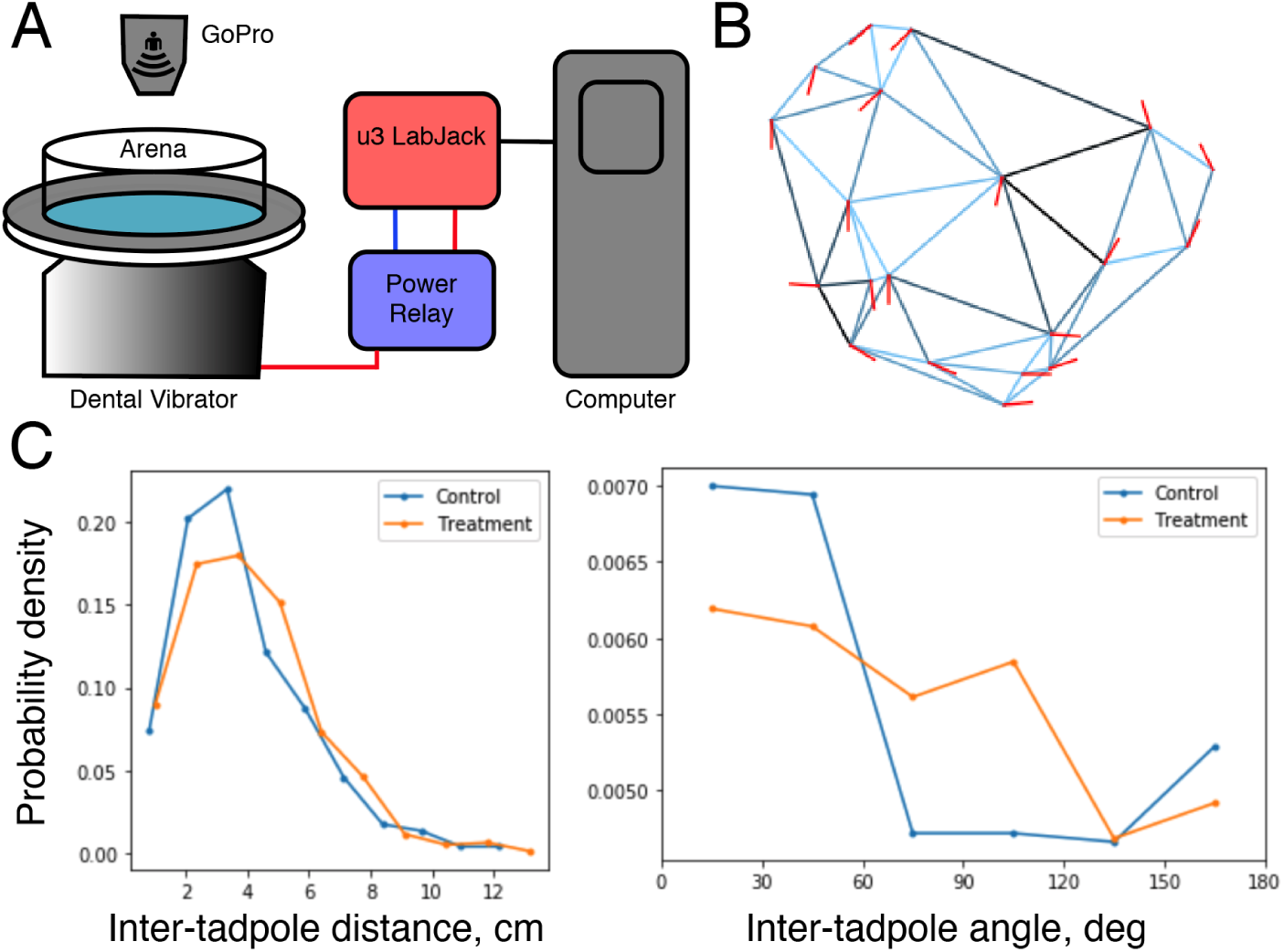
Schematic of equipment setup and analysis. **(A)** Diagram outlining experimental setup using the u3 LabJack, power relay, dental vibrator and GoPro camera. **(B)** Result of triangulation function showing tadpole positions and swimming angle. **(C)** Output of analysis function showing distribution of inter tadpole distances and angles, comparing a control and treatment group.

1. Open the first image in FIJI.
2. Use the line tool to measure the known distance included at the beginning of the experiment.
3. Select `Analyze>Set` scale and include the measured distance in the ‘known distance’ box. Be sure to change the unit of length to the unit used for the measured distance (e.g., cm) and select the ‘global’ box to apply this scale to all subsequent images.
4. Use the multipoint tool to mark the head and gut of each individual tadpole such that each tadpole has a sequential odd number and even number marking for its head and gut, respectively. Care should be taken that the two spots create a vector that aligns with the direction the tadpole body is pointing to.
5. Select `Analyze>Measure` and copy the results into an excel sheet. Repeat this process for every image, adding the measured data to the same excel sheet. When finished, the excel sheet will have 12 blocks of data, representing x-y coordinates for each tadpole, in all 12 images. *Note:* Alternatively, the experimenter may use the ROI Manager tool in Fiji to select the points; this will allow user to save the ROIs with the image.
6. Save the excel sheet as a CSV file `input_control.csv`

### Data analysis – Jupyter notebook

This section describes how to calculate and analyze inter-tadpole distances and relative angles, generating data output files.

1. Run the Jupyter notebooks environment, and navigate to your repository. Open the schooling_analysis notebook.
2. Look through the notebook (perhaps using the “Find” command), and make sure the names of all input and output CSV files are correct. By default, the notebook attempts to read files `input_control.csv` and `input_treatment.csv` from the data subfolder, and then saves the analysis results as `output_processed_control.csv` and `output_processed_treatment.csv` respectively, in the same subfolder.
3. Run the notebook. The program will perform a Delaunay triangulation (a standard way to identify a set of “neighboring” points for every point on a plane), find inter-tadpole distances, and angles between orientations of neighboring tadpoles (**Fig. 1B**). The results, including figures and Kolmogorov-Smirnov test p-values for distance and angle comparisons will appear under respective sections (**Fig. 1C**). Note: While Delaunay triangulation may look somewhat fancy, it is in fact a standard, go-to approach to defining immediate neighbors for a set of points on a 2D plane. In a way, it may be considered a “default” method, and we chose it for the reasons of parsimony. For the analysis of angles, we only compare the orientation of neighboring tadpoles, in the triangulation sense, as there are no reasons to believe that neighboring tadpoles affect each other more strongly, compared to tadpoles that are separated from each other in a school. A reasonable alternative to this approach would be to define an “influence radius”, and for each tadpole, consider all other tadpoles that fall within this radius. However we think that the triangulation approach is more parsimonious, as it is “parameterless”, and we make no assumptions about what this “influence radius” could be. It is also nice (again, in terms of clarity and parsimony) to use the same definition of neighboring tadpoles for the analysis of inter-tadpole distances, and inter-tadpole angles. The simplicity of this method leave less space for fudging the data. In schooling tadpoles, one would observe more short distances, and fewer medium distances, compared to non-schooling (randomly distributed) tadpoles. The distribution of angles will be uniform (flat) for randomly oriented tadpoles, and woudl be declining for tadpoles that co-orient.
4. The last few sections of the code compare data from a single set of experiments (control experiments by default) to reshuffled data, as a way to estimate p-values for a null-hypothesis of “no schooling”. The data is shuffled by taking all observed tadpoles and assigning them to random frames, which preserves the spatial and angular distributions of the data, but removes any patterns, and any coordination between neighboring tadpoles.

### Power analysis

To help with the interpretation of schooling data, we performed rough power analysis (see the notebook `schooling_power_analysis.ipynb` in the Github repository). For this analysis, we generated swarms of “virtual tadpoles” with different amount of schooling (different probability of going in-school, or out of school), and different precision of mutual alignment. In-school tadpoles were placed in the “virtual dish” using a Bridson’s algorithm for Poisson-disc sampling, adjacent to one of existing tadpoles; out-of school tadpoles were placed randomly (uniformly in space). The orientation of tadpoles was set to be in-between a tangent to the radius of the dish (as that is how tadpoles typically swim in a large round bowl) and circularly uniform random orientation, with a mixing coefficient changing from “full alignment” (no noise), to “no alignment” (random orientation).

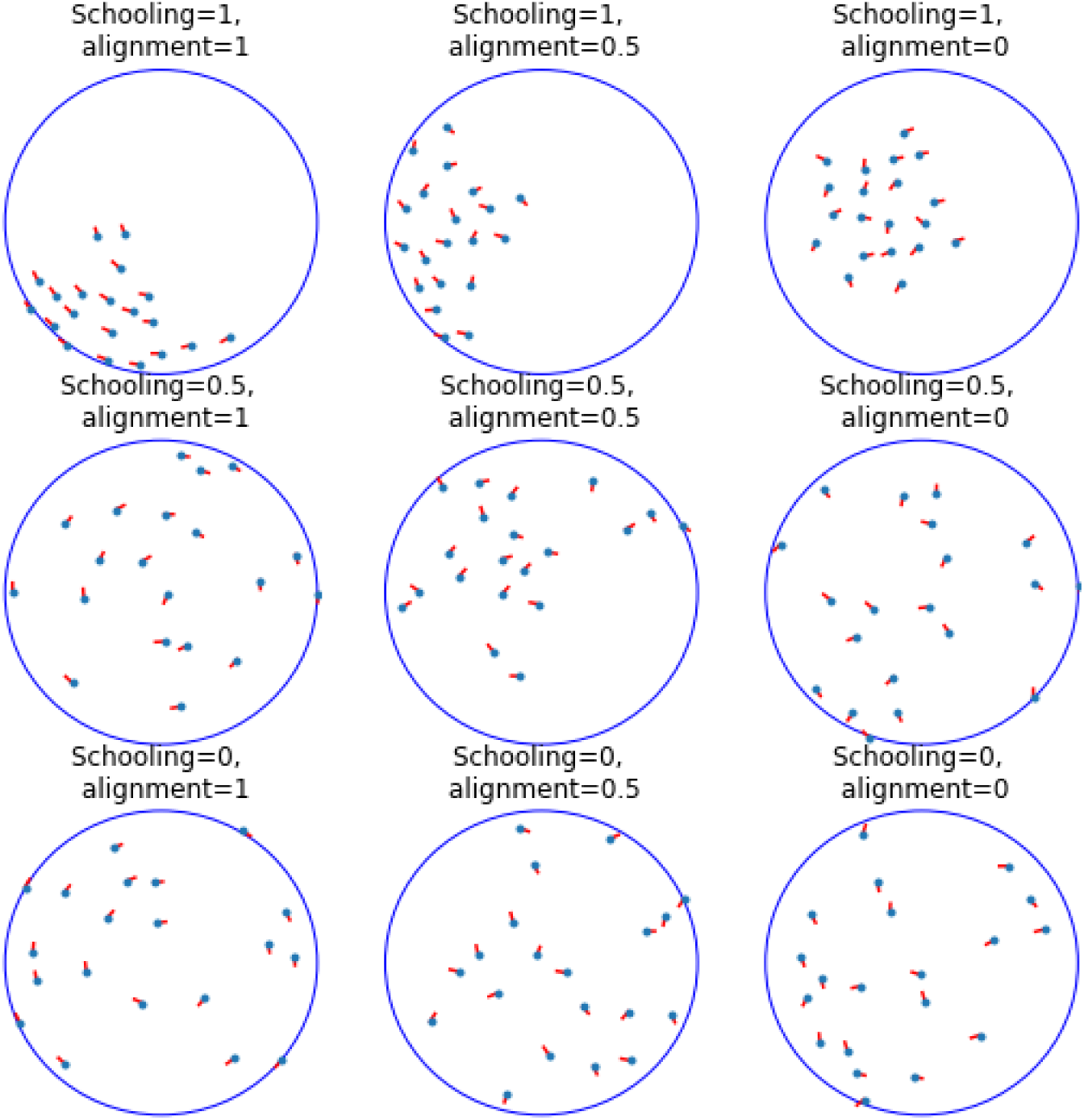

If we process a sample of schooling and non-schooling simulated tadpoles using exactly same routines as we used for real data, we see that in schooling tadpoles, we mostly observe small inter-tadpole distances, and the distribution of distances resembles exponential, while for non-schooling tadpoles we get a flatter distribution, reminiscent of a chi-square distribution. Note that these curves look roughly similar to real biological data presented in the paper.

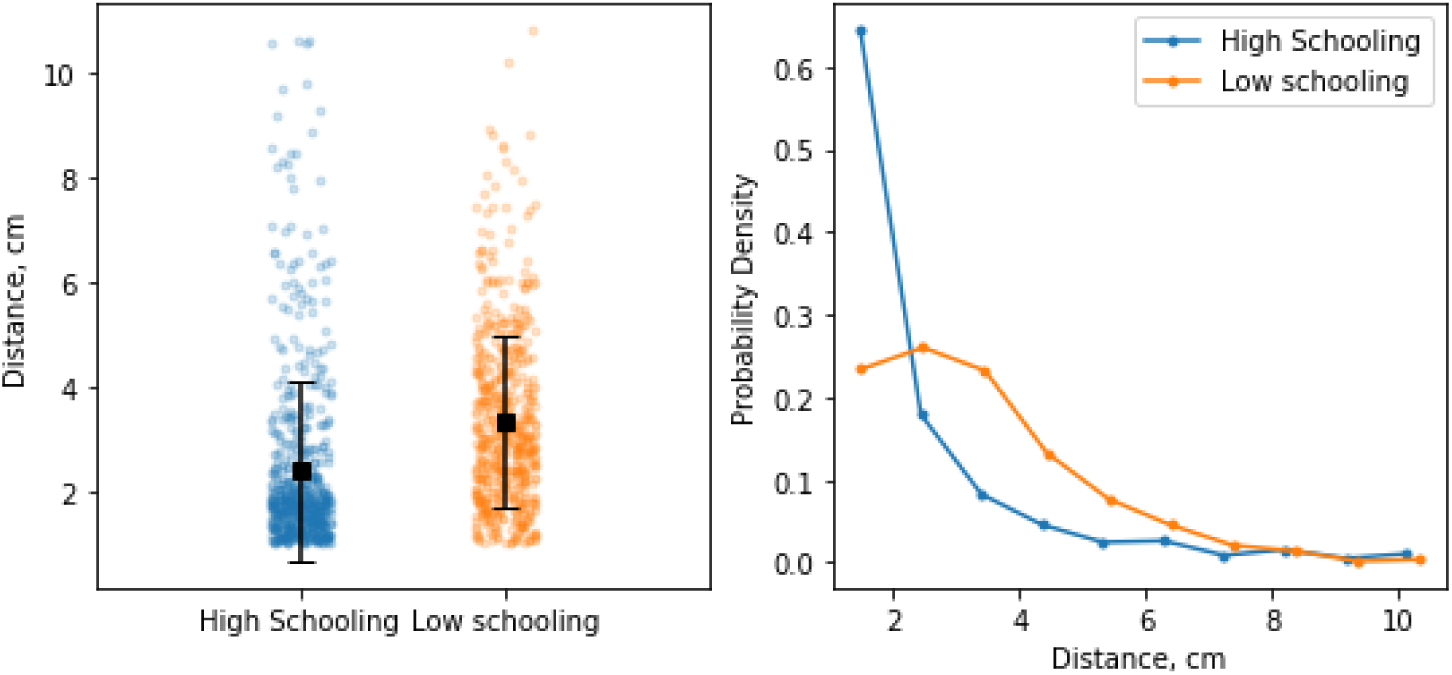

To estimate the power of our tests (their ability to detect a true effect), we cannot rely on the analytical guarantees for Student and Kolmogorov-Smirnov tests, as technically the individual measurements we work with are not independent. This is true for any spatial analysis, both because we look at pairwise comparisons; because all tadpoles share the same limited space, and because consecutive frames are obtained from the same group of tadpoles, just redistributed. One conceptual way to think about it is to assume that we work with a lower, “adjusted” sample size, that is smaller than the “nominal” sample size.

We therefore had to estimate the power of our tests through modeling, by repeatedly simulating 12 “snapshots” of 20 “virtual tadpoles” (the same numbers, as used in real experiments). In each simulation, we kept one “reference” set of “virtual tadpoles” at intermediate values for both schooling (0.5) and alignment (0.5), and let the other set of “virtual tadpoles” have both schooling and alignment change from 0 (no schooling, no alignment) to 1 (perfect schooling and alignment). As we can see from the results below, the angle co-orientation test (green) is incredibly sensitive, and jumps to >95% power even for 5% lower or higher amount of noise in tadpole orientations. For tests on distances, both the t-test (blue), and KS-tests (orange), are less sensitive, and can only detect big differences in schooling. In our simulated example, reference tadpoles joined a school with probability of 0.5 (or otherwise were placed randomly). Compared to this baseline, our tests can detect with ∼80% power a group of tadpoles that joins a school with probability of 0.3, or with a probability of 0.8, which would correspond to a big change in behavior. Nevertheless, the deficits of schooling in real tadpoles are often quite profound, perhaps because schooling is an emergent behavior that relies on several relatively complex and demanding elementary behaviors (swimming, optomotor-response, collision avoidance), and so our methods have a good chance to detect these deficits.

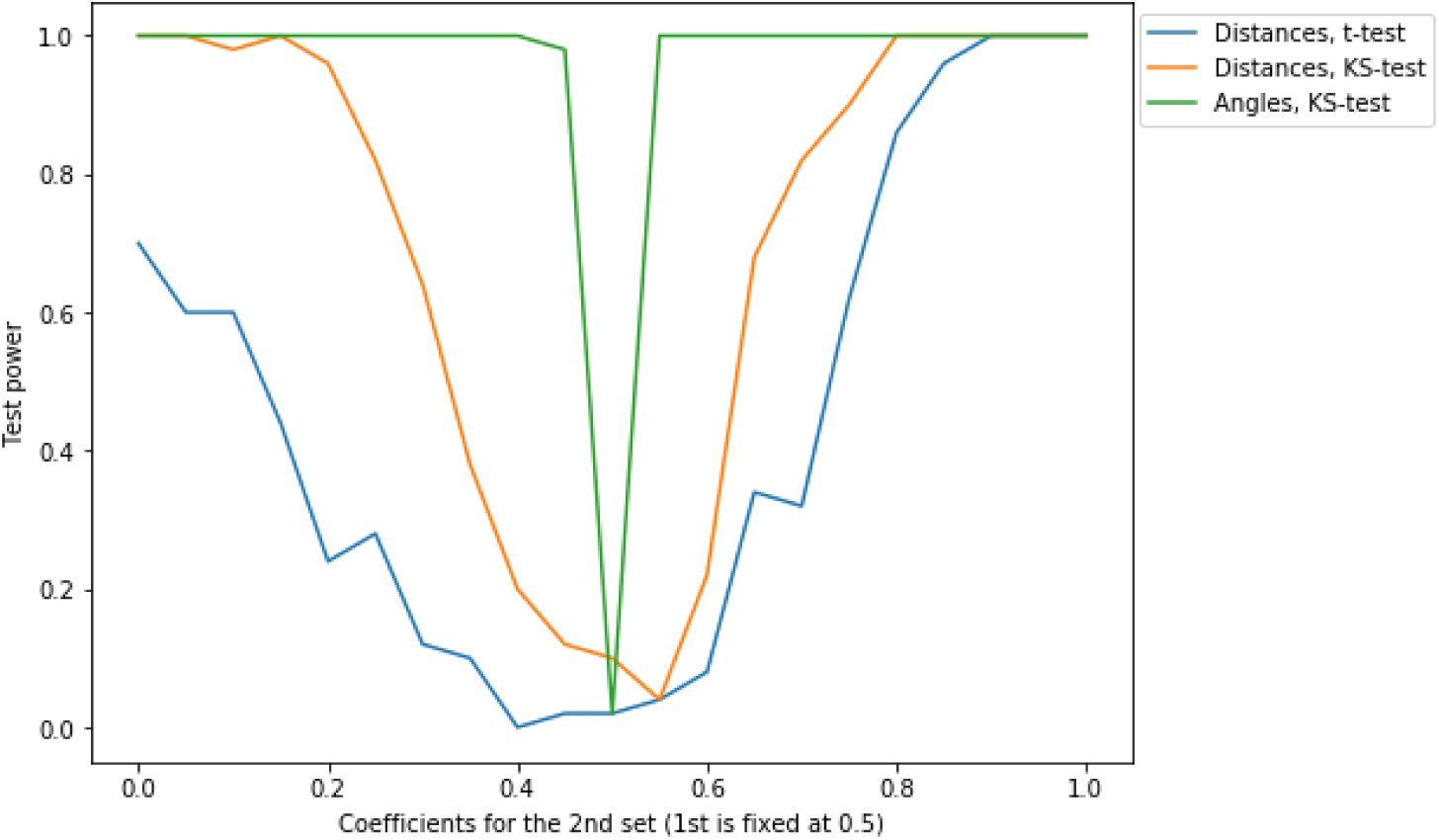

### Troubleshooting

This experiment can be customized a number of ways in order to suit the needs of the experimenter. The schooling_experiment.py script is written to run on a set timer, which can be shortened or lengthened in order to yield more or fewer images for analysis. In order to do this, simply change the value given on line 29 of the code, which is currently set to 4000 seconds by default. Additionally, we have found that tadpole activity decreases as the day goes on, with the highest amount of swimming activity occurring within 1-3 hours into the start of their diurnal cycle (tadpoles are housed in a 12:12 light/dark cycle). As such, care should be taken with regards to when schooling experiments are conducted. Ideally, experiments should be run around the same point of their L/D cycle. Specific criteria should be carefully set up for inclusion of control and experimental groups in the analysis. These criteria could involve a baseline swimming threshold, or accurate schooling in control group. These criteria may vary by age or nature of the experiment, but experimenters should take care to set these a priori criteria for inclusion into their experimental groups.

## Discussion

In our experimental setup, tadpoles organize themselves into small groups within a dish swimming together in one direction. Formation of these schools takes about 30-60 seconds after a vibration stimulus is provided to induce swimming behavior. Thus, we deliver this stimulus with a given time interval so we can sample several schooling configurations from a single group of tadpoles. The included analysis script provides an output file that contains the result of a triangulation used to calculate distances between neighboring tadpoles in each image. It also calculates the relative angles between the orientations of neighboring tadpoles. For normal schooling, the distribution of inter-tadpole distances will be non-random, with more values at shorter distances and longer distances, and fewer “intermediate distances”, representing the tadpole-tadpole distances within a school, and between schools.

Our general approach to experiment design is to have a treatment group and a control group of 20 tadpoles each from the same clutch to account for variations within clutches. Typically, each experiment is run with at least 5 control groups and 5 treatment groups, generally more. Distributions of controls and treatment groups can be compared directly using non-parametric statistics (eg. Kolmogorov-Smirnov test), or each group can be compared to a random distribution to test for presence or absence of schooling (see a simulation notebook on GitHub for more details, and power analysis for these tests). It is important to consider that treatments that result in hyper or hypo activity will almost always show up as differences in schooling behavior. Therefore, it is helpful to assess general swimming activity separately by using video tracking to compare baseline swimming speed between groups.

## 2. Visual Collision Avoidance

In teaching, the best exam questions are those that seem simple at first, but can lead to deep and nuanced conversations. Similarly, to probe brain development, we should look for behaviors that are easy to evoke and quantify, but that are demanding, malleable, and inherently variable. Visual collision avoidance is an example of such a behavior: it is ecologically relevant, robust, and easy to record, but also nuanced, and shaped by the sensory history of the animal. Here we describe how to set up a visual avoidance assay, and how to use it to test sensory processing and sensorimotor transformations in the vertebrate brain.

### Materials

#### Reagents

Tadpole rearing media: 15 mM NaCl, 0.5 mM KCl, 1.0 mM MgSO4, 150 μM KH2PO4, 50 μM NaHPO4, 1.0 mM CaCl2, 0.7 mM NaHCO3, 0.5 mg/L methylene blue, in DI H2O; pH 7.2-7.6. Other rearing media, such as the popular “0.1X Steinberg solution”, are also acceptable.

#### Equipment

- Projection table Use 1100 cm (36’) of 20×40 mm (1’’×2’’) pine beam; 32 wood screws (e.g. #8 x 1’’ flat head); 4 furniture felt pads; 2 pieces of clean acrylic 31×31 cm (1’×1’), and a white disposable plastic apron (Fig 1A).
- Short throw projector (one capable of producing a focused image at a distance of ∼1 m), e.g. AAXA P300 As an alternative, use a CRT monitor, or an iPad-like tablet.
  - Computer with Internet access
  - Plastic Petri dish (8.5 cm in diameter)
  - Web-camera

**Optional:**

- Cardboard box to screen the arena
- IR light source (e.g. Univivi U48R LED wide angle source for CCTV security)
- Acrylic light filters
- Tracking software, such as:

http://ctrax.sourceforge.net

http://sourceforge.net/projects/buridan/

## Methods

### Projecting device

As the visual stimulus is projected on the floor of the chamber, the tadpole should be able to see the image even at sharp angles. We therefore cannot use a standard LCD monitor for visual stimulation, but have to use either a projector, an old CRT monitor placed horizontally, or a tablet. As a test, if you cannot see the image on the screen when looking along the surface, the tadpole will not see it as well.

We recommend to use a short-throw USB pico-projector, and a screen made of a piece of white disposable polyethilene apron, fixed between two pieces of clean acrylic (**Fig 2A**). The downsides of CRT monitors is that the surface is never quite flat; they heat the water; and the video recording is compromised by a refresh beam artifact. The protocol also works with high-end tablets, but we have not explored this approach systematically.

**Figure 2.**
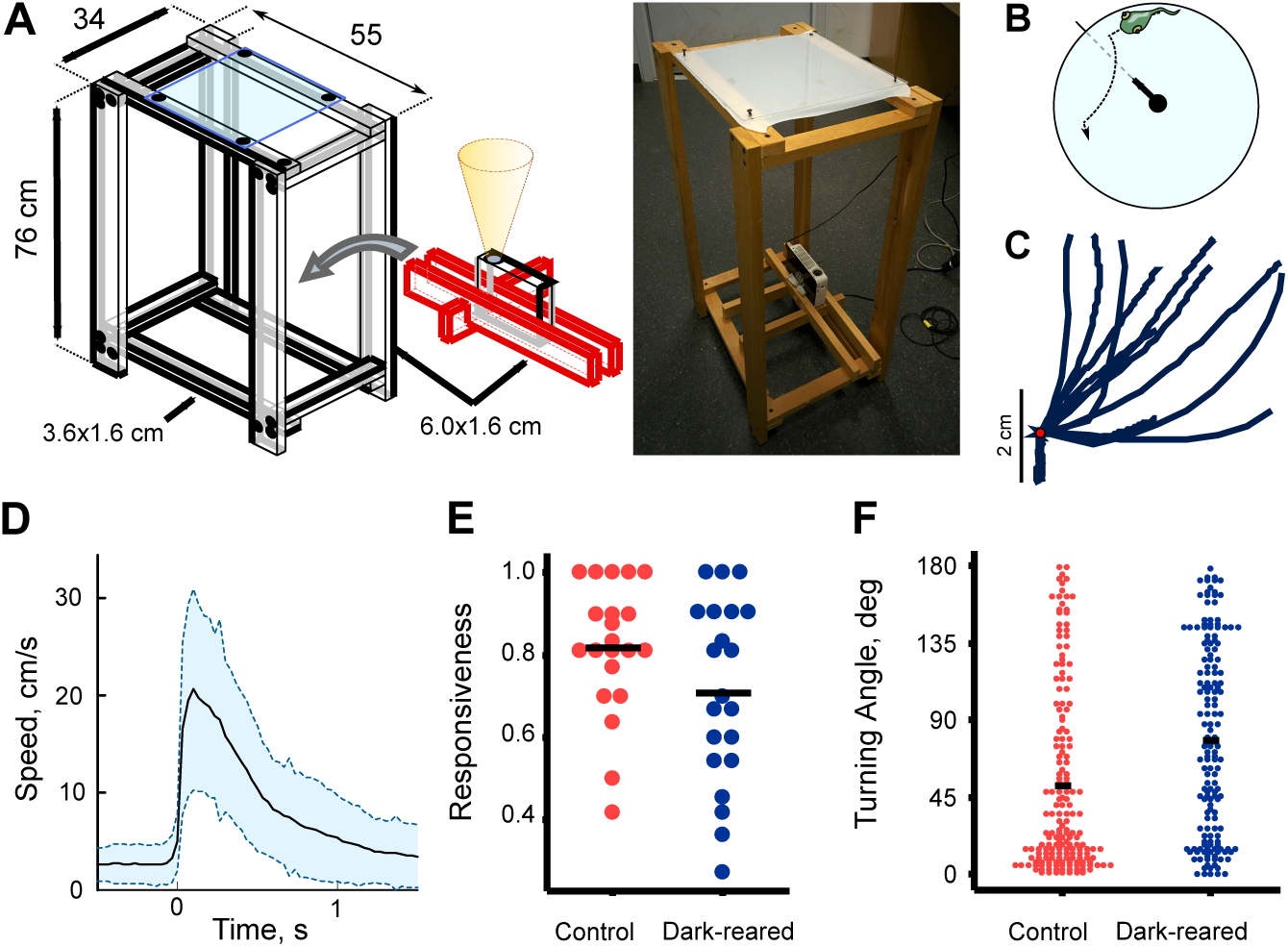
**A**. The projection table. **B**. Schematics of the experiment. **C**. Representative avoidance trajectories. **D**. Average (solid) and standard deviation (dashed) values of swimming speed for 500 fast avoidance responses. **E**. Response probabilities for normal and dark-reared tadpoles. **F**. Turning angles for normal and dark-reared tadpoles. This data was previously presented in (Khakhalin at al. 2014) and (Ramirez-Vizcarrondo et al. 2015), but was analyzed in a new way for this paper.

#### Recording

1. In a browser, open the repository of Xenopus behavioral protocols: https://github.com/khakhalin/Xenopus-Behavior Navigate to the current version of the Collision Avoidance Stimulator (**Fig 2B**). Project the image.
2. Adjust the background lightness, to keep the contrast high without blinding the tadpole. To assess the optimal lightness for your setup, run a series of experiments with different contrasts, and pick the contrast with the highest response rate.
3. Place a Petri dish on top of the screen, and fill it with tadpole rearing media, 1-1.5 cm deep. Make the arena of the program match the position of the Petri dish; adjust the radius if needed.
4. Set the stimulation parameters. Wait for at least 20, better 30 s between the stimuli, to prevent habituation. Make the black dot ∼5 mm in diameter (comparable to the size of a tadpole), traveling at a speed of 3-5 cm/s (comparable to the speed of a tadpole). Making the circle faster or smaller (e.g. v=4 cm/s, d=4 mm) increases the chances of triggering a “fast” poorly coordinated escape response, while keeping the circle slower or larger (e.g. v=4 cm/s, d=12 mm) allows tadpoles to implement more spatially informed course corrections (Khakhalin et al. 2014; Khakhalin 2019).
5. Target the circle using LEFT / RIGHT keys, then send it towards the tadpole by pressing the UP key. You can return the circle to the center by hitting the DOWN key. Sending the circle towards the tadpole sets the timer in the left bottom corner of the screen. The timer controls the color of the circle, making it pale during the inter-stimulus interval, to prevent habituation.
6. Before running actual experiments, practice your targeting. As healthy tadpoles tend to travel around the edges of the arena, it is usually enough to keep the target fixed, and time your sending the circle on the collision trajectory. Do not press the LEFT / RIGHT keys while the circle is in motion, as lateral motion increases the speed of the stimulus.
7. During actual experiments, place tadpoles for each treatment in a separate bowl, and make the experiment alternate between groups, while keeping them blinded to the identity of each group.
8. Record a video for post-processing, but also quantify each trial as you go, as either a “success “(if the avoidance response was triggered; **Fig 2C, D**) or a “failure”.

### Analysis

The simplest experimental design involves comparing rates of avoidance responses across different treatment groups (**Fig 2E**). This analysis requires minimal equipment, can be done by eye during the experiment, and later verified from the video (scored by another person). This design is also appropriate for teaching labs. To better probe brain development, one can sweep across a range of values for circle speeds, sizes, or contrasts, and compare the results between groups (Henriet et al. 2017).

A more sophisticated analysis involves tracking the tadpole position at every frame, which can be done manualy (ImageJ), or automatically, using commercial (Noldus EthoVision; CleverSys AquaScan) or open source software (CeTrAn, C-trax, etc.; Colomb et al. 2012; Chao et al. 2015). Tracking of both the stimulus and the tadpole gives access to several more variables, such as the peak swimming speed, the angle of turn during the avoidance maneuver (**Fig 2F**), and the distance to the circle when the response is triggered. It also allows automated classification of responses into “successes” and “failures”.

As not all software solutions can track both the circle and the tadpole, another option is to add a near-IR light source below the projection screen, and turn the camera to IR-only, by replacing a glass IR filter inside it with a stack of of three colored (red, green, and blue) acrylic filters (Truszkowski et al. 2017). In this configuration, the camera would only record the tadpole, but not the stimulus, which still allows for some analysis of trajectories.

The stimulation program presented here can also be used for multisensory experiments, as it can deliver a sound either when the circle hits the wall, or immediately before, or immediately after that. This protocol however falls out of scope of this paper.

### Troubleshooting

**Problem:** Tadpoles are sluggish and do not swim, staying in one place; yet the moment they are returned to the tank, they start swimming again.

**Solutions:**

- Give the tadpole a minute to acclimate, then startle it with a gentle tap. Ideally, a tadpole should always be swimming, with a speed of 1-3 cm/s.
- Make sure the media is cold enough. *X. laevis* tadpoles prefer temperatures slightly colder than room temperature (18° to 21°C). If the projector heats the dish, replace the media regularly.
- Add ambient light to the chamber: e.g. make the screen around it white. Tadpoles stop swimming if the environment is too dark.
- Run the experiments earlier in the morning: on a 12/12 light cycle with “dawn” at 7 am, tadpoles seem most active between 9 am and 12 pm, and often stop responding in the afternoon.
- Do not feed tadpoles before the experiment.
- Adjust the brightness down to reduce retinal bleaching, or increase the contrast.
- If working without a screen, reduce movement, to avoid visual habituation.

## Discussion

A typical fast avoidance maneuver involves a sharp turn by ∼90±40° performed at a distance of ∼1 cm from the circle, followed by a brief acceleration to 12±4 cm/s. A slow course correction is produced in response to larger, slower circles, and involves a shallower turn by ∼70±50° and acceleration to ∼7±4 cm/s (Khakhalin et al. 2014).

A typical responsiveness for this protocol is 80% for control Nieuwkoop-Faber stage 49 tadpoles (15-30 days post-fertilization, if raised at 20°C; Khakhalin et al. 2014), but it goes down to 70% in dark-reared tadpoles (Ramirez-Vizcarrondo et al. 2015), 50% for tadpoles adjusted to strong visual stimulation (Jang at al. 2016); 40% for animals with mild neurodevelopmental abnormalities (James et al. 2015), and 20% in nutritionally restricted tadpoles raised in isolated wells (Khakhalin, unpublished). A power analysis shows that with 20 stimuli per tadpole (10 minutes of recording), base response probability of 70%, 20 animals in each group, and α = 0.05, one can detect a drop of responsiveness to 60% with 80% power. Together, this makes visual collision avoidance a powerful tool to dissect abnormalities in the sensorimotor system of Xenopus tadpoles, from retinal disfunction, through sensorimotor transformation proper, and down to locomotor deficits.

## Appendix A. Reading Media

### 10% Steinberg’s Rearing Solution

The recipe below describes how to prepare **10X Stock** Steinberg’s solution. In practice, tadpoles are reared in 10% Steinberg’s solution. It means that the recipe below gives you a fluid that needs to be diluted 100 times to produce appropriate Xenopus rearing media. The reasons for this confusing naming situation are purely historical.

| Reagent | Amount |
| --- | --- |
| NaCl | 34.0 g |
| HEPES | 119.0 g |
| Ca(NO<sub>3</sub>)<sub>2</sub> · 4H<sub>2</sub>O | 0.8 g |
| MgSO<sub>4</sub> · 7H<sub>2</sub>O | 2.06 g |
| KCl | 0.5 g |
| dH<sub>2</sub>O | 1 L |

Preparation protocol:

1. Completely dissolve all reagents in 850 mL of dH2O.
2. Adjust pH to 7.5 with 10N NaOH. To prepare 10N NaOH, slowly dissolve 40g NaOH in 100 ml dH20. Store at room temperature.
3. Add dH2O to reach final volume of 1 L and store at 4°C.
4. Dilute 1:100 to create 10% Stenberg’s, to use as rearing solution for tadpoles.

